# In vivo time-lapse imaging and analysis of mitochondria in neural progenitor cells in the developing brain

**DOI:** 10.1101/2023.07.31.547741

**Authors:** Martin S. Feng, Maggie R. Kettelberger, Jennifer E. Bestman

## Abstract

Neural progenitor cells (NPCs) are the highly polarized dividing stem cells of the developing brain that give rise to all neurons and glia. Early on, NPCs divide symmetrically and expand the pool of progenitor cells, but as development continues the NPCs begin to asymmetrically divide to produce neurons. The mechanisms that govern this irreversible commitment to neurogenesis are not fully understood, but in other stem cell populations the regulation of mitochondria and cell metabolism is key to controlling stem cell fate. Here we use timelapse 3D confocal microscopy to observe NPCs, their cellular progeny, and their mitochondria in the developing *Xenopus* tectum. Our results track individual NPCs over days and show that they contain abundant mitochondria that form complicated networks distributed throughout the cells. We find that NPCs preparing to divide shift mitochondria toward the cell body where they become asymmetrically distributed, suggesting that the cells control which progeny inherit mitochondria. This uneven distribution of mitochondria in cell preparing to divide led us to test the role that mitochondria play in cell division. We overexpressed the mitochondrial biogenesis master regulator, PGC-1a, which induced the NPCs to asymmetrically divide and produce neurons, while PGC-1a knockdown limited neurogenesis. Together these data suggest that the regulation of mitochondria by NPCs prior to cell division and their unequal inheritance during cell division, contributes to the fate of the newborn cells in the developing brain.

## INTRODUCTION

Nearly every eukaryotic cell is beholden to its mitochondria, the multifunctional, genome-containing organelles that not only generate ATP, but also regulate Ca^++^ and serve as hubs to many biosynthetic pathways (Monzel et al., 2023). In most cells, mitochondria form dynamic, branching networks (Collins and Bootman, 2003) and through the process of fusion and fission mitochondria proliferate and rearrange as cellular demands change (Chan, 2020). Since cells are incapable of generating mitochondria *de novo*, they must inherit them from their parent during cell division. The complex morphology of mitochondria is an obstacle to mitosis, as the networks must be broken apart prior to cell division. Indeed, mutations in mitochondrial dynamin (Drp1) that prevent mitochondrial fission and the disassembly of the network directly alter cyclin E protein levels (Mitra et al., 2009; Parker et al., 2015), and disrupt progress through the cell cycle (Taguchi et al., 2007).

How mitochondria and mitochondrial networks are divided in preparation for cell division is associated with the ability of cells to proliferate (their “stemness”), and the fate of their nascent cell progeny (Kumar Sharma et al., 2022). No clearer example exists than oogenesis, during which time the maturing oocytes preferentially inherit mitochondria compared to their polar body siblings (Dalton and Carroll, 2013). When mitochondria segregation is inhibited during oocyte maturation, polar bodies fail to extrude, resulting in dysfunctional oocyte formation (Lee et al., 2022). Similarly, failures in the asymmetric inheritance of healthier mitochondria into the bud of dividing *Saccharomyces* yeast halt the cell cycle and produce cytokinesis failure (Pernice et al., 2017).

Stem cells actively regulate mitochondrial structural dynamics and mitochondrial inheritance. Therefore, it is surprising that across tissues and organisms stem cells do not require or, indeed, actively inhibit gene expression and metabolic processes that require mitochondria (Angelopoulos et al., 2022; Khacho et al., 2019; Petridi et al., 2022). For brain stem cells, the reliance on non-mitochondrial ATP production was first shown *in vivo* by Agathocleous et al. (2012) who found that despite having mitochondria and oxygen, neural progenitor cells of *Xenopus laevis* retina preferentially used glycolysis, a non-mitochondrial mechanism for generating ATP. Moreover, when glycolysis was inhibited, cell division stalled and the progenitor cells died (Agathocleous et al., 2012). As stem cells differentiate and their pluripotency declines, mitochondria network complexity increases and ATP production through mitochondrial oxidative phosphorylation substitutes for glycolysis (Bahat and Gross, 2019).

In the developing brain, the dividing stem cells that give rise to neurons and glia are radial glial neural progenitor cells (henceforth, NPCs;(Sild and Ruthazer, 2011). NPCs are unusual cells in the brain because their cell bodies (or short apical processes) are in direct contact with the brain’s ventricular surface, and the cells also make contact to the brain’s outer, pial (basal) surface with an enlarged endfoot. Spanning these two cell compartments is a slender radial process (the basal process (Bystron et al., 2008) that grows hundreds of microns as the brain expands in size during development. This unique morphology grants NPCs access to signals arising from all layers of the developing neural tissue around them, the vasculature, and the cerebral spinal fluid. These inputs form the neurogenic niche which is essential in directing the fate of the NPCs (Bjornsson et al., 2015). Cell division for such polarized cells presents a logistical problem: even when NPCs divide symmetrically to generate two new NPCs, cellular contents are rarely equally divided (Kosodo et al., 2008), and typically one cell begins its life essentially as a bud that must rebuild the NPC morphology and its sibling inherits the bulk of the cellular components of the parent (the radial process and pial endfoot (Miyata et al., 2001).

Foundational electron microscopic reconstructions of radial glial cells in the developing cortex along with more recent studies have reported that mitochondria are found throughout all compartments of the NPCs (Rakic, 1972; Rash et al., 2018). Careful examinations of the morphology of the mitochondria in the cell bodies of dividing cortical stem cells demonstrated that they are relatively complex but become fragmented within the cells committed to differentiate into neurons (Iwata et al., 2020; Khacho et al., 2017). When mitochondrial networks are prevented from disassembling through the manipulation of the mitochondrial fission factor Drp1, the ability of cortical NPCs to form neurospheres in vitro is limited, indicating that mitochondrial dynamics play a role in proliferation (Khacho et al., 2016). Recently, Iwata et al. (2020) showed that similar manipulations of mitochondrial fission and fusion conducted soon after mitosis were also able to prevent newborn cells from committing to a neurogenic fate (Iwata et al., 2020). Furthermore, once committed to a neuronal fate, continued mitochondrial structural dynamics supports the energic needs of neurons and is critical for their maturation and survival (Ishihara et al., 2009; Steib et al., 2014; Wakabayashi et al., 2009). These data demonstrate that mitochondrial dynamics regulate the fate of NPC progeny, but leave open questions about their role the NPC parent cell as it prepares to divide.

NPCs in the *Xenopus laevis* tectum (in mammals, the superior colliculus), are the stem cells that generate neurons and glia in this major visual input processing area of the brain. Tectal NPCs, like their cortical counterparts, express key stem cell markers such as Sox2 and the microfilament protein, vimentin (Bestman et al., 2015; Sharma and Cline, 2010). The maturation of the visual circuitry is accompanied by steady cell proliferation that continues throughout visual system development (Lazar, 1973; Peunova et al., 2001; Straznicky and Gaze, 1972) even after the visual system is functional (Dong et al., 2009).Tectal NPCs are a mixture of symmetrically dividing cells that divide and expand the number of progenitors, and those that have begun to asymmetrically divide and generate neurons (Bestman et al., 2012; Herrgen and Akerman, 2016). Akin to how exercise and cognitive activity alter adult neurogenesis in mammals (Kempermann, 2019), levels of cell proliferation in the tectum are plastic, and neurogenesis increases or decreases in response to the animal’s visual experience (Bestman et al., 2012; Sharma and Cline, 2010). Because *Xenopus* tadpoles are transparent and their brain lies just beneath their skin, experiments designed to reveal the molecular mechanisms of neurodevelopment at resolution of molecules to neural circuits are ideal for this system (Pratt and Khakhalin, 2013).

Here we use timelapse 3D confocal microscopy to observe NPCs, their cellular progeny, and their mitochondria in the developing *Xenopus* tectum. The unparalleled advantage to this longitudinal imaging approach is that the proliferation of individual cells can be observed over days, but the brain environment (the “neurogenic niche” of the brain) that is critically important for controlling cell proliferation (Bjornsson et al.; Cairns et al., 2011), is completely intact. Our results show that tectal NPCs contain abundant mitochondria that form complicated networks distributed throughout the cells. After imaging individual NPCs over multiple days, our retrospective quantitative analysis of the fluorescently-tagged mitochondria revealed that 24 hours prior to cell division, mitochondria in the radial process shift toward the cell body of the NPCs. This bias in the distribution does not occur in NPCs that remain quiescent and do not divide. Within the cell body, mitochondria are not distributed equally prior to cell division, suggesting that that the parent NPCs are preparing for the asymmetric inheritance of mitochondria by their progeny.

To test the importance of mitochondria to NPC fate, we manipulated mitochondrial biogenesis by overexpressing or knocking down the “master regulator” transcription factor, peroxisome proliferator-activated receptor gamma co-activator alpha (ppargc1a) which encodes PGC-1a (Handschin and Spiegelman, 2006; Scarpulla, 2011). PGC-1a overexpression in neurons has been shown to be significantly increase mitochondrial biogenesis, induce the expression of electron transport chain proteins, and improve mitochondrial metabolic pathways so effectively it is being investigated as a possible therapeutic target for the treatment of neurodegenerative diseases (Rosenkranz et al., 2021). In the mouse brain, PGC-1a loss causes neurodegeneration (Lin et al., 2004) and in humans, loss of PGC-1a is associated with a number of neurodegenerative diseases (Lucas et al., 2014). We found that NPCs overexpressing PGC-1a were significantly more likely to generate neurons while knocking down ppargc1a expression interfered with neurogenesis. These data suggest that regulation of mitochondrial biogenesis prior to cell division is contributes to fate of the NPC progeny.

## Methods

### Tadpole rearing and tectal cell transfection

Albino *Xenopus laevis* tadpoles were purchased from *Xenopus* express (*Xenopus*.com) as embryos and raised in Steinberg’s solution (58.0 mM NaCl, 0.67 mM KCl, 0.34 mM Ca(NO3)2, 0.83 mM MgSO4, 3.0 mM HEPES, pH 7.2) in12 hour light/12 hour dark cycle incubators at 22-23° C until stage 46 (Nieuwkoop and Faber, 1994). Tadpoles were fed *Xenopus* express Premium Tadpole Food beginning at stage 46. After the first confocal image, the tadpoles were housed singly in 6-well cell culture plates. The animal protocol was approved by the Institutional Animal Care and Use Committee of William & Mary in accordance with NIH guidelines.

### Tectal cell transfection and molecular reagents

Cells in the tectum were transfected using in vivo electroporation of stage 46 animals anesthetized in 0.01%MS-222 (3-aminobenzoic acid ethyl ester, Sigma) dissolved in Steinberg’s media. 100-500ng/µl plasmid solutions mixed with ±0.05% fast green dye were injected into the midbrain ventricle and 3 or 4, 40V pulses were delivered to the midbrain using platinum electrodes using a Grass SD9 stimulator (Haas et al.). The plasmids used in this study are modified from the pSox2-bd::FP construct (Addgene plasmid #34703; Bestman et al., 2015), a plasmid that contains a weak FGF4 promotor regulated by 6 repeats of the Sox2 transcription factor binding domain. This regulatory domain drives the expression of Gal4-VP16 and so fluorescent protein expression is in turn driven by upstream activating (UAS) sequences (Bestman et al., 2015). Because Sox2 is expressed only in proliferating cells, this regulatory cassette preferentially drives expression in tectal NPCs compared to differentiated neurons which do not express Sox2 at high levels.

Plasmids were constructed using Gateway Cloning (Invitrogen). We used pSox2-bd::FP (Addgene plasmid #34703) to express EGFP in our cells and this plasmid was used to create p5E-Sox2 (L4-R1), a 5’ element Entry plasmid that contains the Sox2-bd regulatory domain including gal4-vp16 from pSox2-bd::FP. Two different cytosolic red fluorescent proteins were used in our experiments, TagRFP-T, a gift from Michael Davidson (Addgene plasmid #58023).

The ppargc1a coding domain (XM_018243070) was cloned from X. *laevis* tadpole (stage 40) cDNA using SuperScript IV reverse transcriptase (Invitrogen) (F: gatccaatcaggggctggat; R: cctacgcaggcttctctgc) and recombined with the gateway pDNOR221 middle entry plasmid to create pME_ppargc1a. To express ppargc1a in NPCs, the p5E-Sox2 (L4-R1) was combined with *Xenopus laevis* ppargc1a separated from tagRFP-T by the porcine teschovirus-1 2A (P2A) “self-cleaving” peptide. In this plasmid, pSox2-bd::ppargc1a_p2a-TagRFP-T, the P2A peptide efficiently separates two proteins during their synthesis (Donnelly et al., 2001) so that the expression of red fluorescence identifies the cells that express an untagged ppargc1a.

To label mitochondria, we made a UAS::mitochondrially-localized EGFP using the repeated UAS sequence derived from the p5E-4xnrUAS (L4-R1), a gift from Marnie Halpern (Addgene plasmid # 61372) with EGFP containing the canonical n-terminal mitochondrial localization sequence from cytochrome c oxidase subunit VIII (mito-EGFP) to create p4XUAS::mitoEGFP. This plasmid could be combined with any that contain the pSox-bd regulator domain to produce coexpression in the NPCs.

To knockdown gene expression we used antisense morpholino oligonucleotides (GeneTools, LLC) designed to target the start site of *Xenopus laevis* ppargc1a mRNA (bp -4 to +21 of XM_018243070.20 and a control morpholino with no targets in the *Xenopus laevis* genome (Fisher et al., 2023). The ppargc1a morpholino sequence is GTTACACATGTCCCACGCCATCCAG and the control morpholino sequence is TAACTCGCATCGTAGATTGACTAAA. The morpholinos were diluted to make 1mM morpholino stock solutions and further diluted to 100 µM and mixed with pSox2-bd::EGFP plasmid for electroporation (Bestman and Cline, 2020).

### In vivo imaging and analysis

Approximately 24 hours after electroporation of the plasmids, tadpoles were anesthetized in 0.01% MS-222 and placed in a custom-built chamber with a coverslip directly on the skin on their heads. Confocal z-stacks of labeled cells in each lobe of the tectum were acquired at ≤1 µm z-step intervals from the dorsal surface to ∼150 µm depths using Slidebook image acquisition software (3i, Inc.) each day for at least 3 days. Between imaging sessions, the tadpoles were returned to a well of a six-well culture plate. We used a VIVO spinning disc confocal system (3i, Inc.) composed of a Yokogawa CSU-X1 spinning disk confocal and a Photometrics Prime 95B sCMOS camera mounted on a Zeiss Axio Examiner microscope equipped with a C-APO 63x/1.15 objective. The 561 nm laser was used to excite TagRFP-T and the 488 nm laser excited EGFP. Signals were distinguished with single bandpass Semrock 617/73 nm and 525/30 nm filters or a dual 525/21-621/58 nm filters. Details of the microscope system are entered here: <u>FPBase</u> (Lambert, 2019). Acquisition settings were adjusted to prevent pixel saturation and identical settings were used throughout an experiment. We selected tadpoles with few spatially-distributed cells for the timelapse image analysis so that the morphology of individual cells could be reconstructed without ambiguity. For the morpholino knockdown experiments, we selected tadpoles with tectal lobes that contained an average of 12 EGFP-positive cells (95% confidence intervals = 9 and 15) on day 1. The same image was acquired on day 2 and day 3.

Image analysis was conducted using the raw, 16 bit image stacks using Slidebook and FIJI/Image J (Schindelin et al., 2012). Although many image analysis tools have been developed to quantify mitochondria (Chu et al., 2022), few are suitable for the large 3D confocal stacks of the tectal NPCs which have a large (∼10-fold) range in mitoEGFP intensities with large variations in mitochondria size and network complexity that fail to be accurately quantified with automated analysis methods. Therefore, we developed methods to quantify mitoEGFP signal without threshold or manual selection so that it could be compared between cell groups. To analyze the distribution of mitochondria in the radial process, Simple Neurite Tracer (SNT; Arshadi et al., 2021) was used to make the 3D reconstruction of each radial process from the soma to the most distal pial endfoot. After subtracting the background, the intensity of the mitoEGFP channel along the radial process was extracted using SNT’s plot profile (max intensity, 3 µm radius), and these position and intensity values were then normalized to their maximum. From these data we calculated the cumulative normalized intensity (CNI) for each cell: the total mitoEGFP signal along the radial process from the cell body to the distal pial endfoot was summed and scaled to 1. If mitochondria are distributed equally, the CNI value at any point along the radial process would equal the unitary slope (e.g., at the 50% position along the process, the CNI would equal 50%). CNI values were calculated at the 10% intervals along the radial process. Data from individual cells are shown along with the average CNI values for the NPC groups.

To create the kymograph of the radial process, any drift of the cell that occurred during the timelapse acquisition was corrected using the descriptor-based series registration tool (Preibisch et al.) to align the images. After reconstructing the radial process with SNT as described above, the SNT path was used to produce a 40 µm straightened view of the radial process. From this image, a 3 µm diameter line was drawn along the radial process and MultiKymograph was used to assemble the kymograph montage.

To measure the distribution of mitochondria in the NPC somata and calculate the magnitude and angle of the mitoEGFP signal for each cell, we used the ROI manager in FIJI/ImageJ to create 24 lines that share an origin point (a wheel with 24 spokes) that radiated out at 15° or 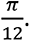 radian intervals. This ROI group was positioned so that the NPC’s radial process emerged at 0° and using the Multi Plot tool, the intensity along the line was measured from the mitoEGFP channel. These values were normalized to the maximum value for each cell before the intensity was summed within each radian segment. The X and Y components for each segment were calculated in order to derive the mitoEGFP vector for the segment, these 24 vectors were used to calculate the total mitoEGFP vector magnitude and angle for each cell. To compare NPC groups, the average magnitudes were calculated, and comparisons were made using Mann-Whitney U tests.

We assessed the percentages of neurons and NPCs in the morpholino-expressing tectal lobes using the ROI manager (FIJI/Image J) to mark and label each cell within the 3D confocal stack and identify it as a neuron or NPC. The image files were coded so that the analysis was done without knowing the identity of the images. NPCs were identified by their morphological features: they had a pial endfoot, simple unbranching radial process and angular cell body apposed to the ventricle. Cells that failed to have these features and/or had branching dendrites assigned as neurons. Cells that were ambiguous were excluded from the analysis.

Figures show maximum z-projections of background subtracted confocal stacks unless indicated that the cell was rotated and projected in another orientation. The 8-bit images were created in FIJI/ImageJ and figures were assembled with Adobe Photoshop and Adobe Illustrator. Because of the large range of mitoEGFP intensities within one cell, to create images that can display the mitochondria with relatively low EGFP signals, the maximum intensity value has been altered in some 2 channel images, but the inset inverted grey scale images show the full dynamic range mitoEGFP signal.

### Statistical Analyses

Statistical analyses were conducted using JMP 17.0 software. Data are reported as means ± standard error of the mean in the text. Box plot figures show the quantiles, means (large lines) and medians with individual data points; p-values < 0.05 are considered significant. The tables report the mean, standard deviation and 95% confidence intervals along with the statistical tests. Pearson’s Chi Square analysis was used to describe differences in the proportions of cell populations. Nonparametric Mann-Whitney U tests were used to compare groups and Wilcoxon Signed Rank tests were used to test paired data.

## RESULTS

### In vivo imaging of tectal neural progenitor cells shows abundant and distributed mitochondria

*Xenopus* neural progenitor cells (NPCs), like all stem cells, express SRY (sex determining region Y)-box 2 (Sox2), a core component of the transcriptional regulation that maintains multipotency in undifferentiated cells (Liu et al., 2013). We capitalize on this defining characteristic of the NPCs to drive cell-specific expression from the plasmid fluorescent protein expression vectors (Bestman, 2012 and see methods). With in vivo electroporation, individual NPCs of stage 46 *Xenopus laevis* tadpoles (Nieuwkoop and Faber, 1994) were transfected with plasmid DNA to label the cells and their mitochondria. After 24 hours, 3D confocal stacks were acquired of the full structure of fluorescently-labeled NPCs in the intact brain each day for at least 3 days. From this timelapse data, we aimed to identify how these very polarized cells divide themselves and their EGFP-labeled mitochondria.

Figure 1 shows a tectal NPC labeled with cytosolic tagRFP-T (magenta) and mitoEGFP (green) captured each day for 3 days. The diagram illustrates the typical position of tectal NPCs in the brain (Fig. 1A). The maximum z-projection of the 75 µm confocal stack of the NPC is oriented so that its ventricularly-apposed soma is at the bottom of the image and its radial process that extends to the dorsolateral surface of the tectum and ends with the pial endfoot is at the top of the image (Fig. 1B). The 90°-rotated X-projection better reveals the radial process projecting dorsolaterally (Fig. 1C). The mitoEGFP signal (in inverted grey scale) of the entire cell (Fig. 1D) reveals that mitochondria are distributed throughout the cell including the NPC’s pial endfoot (Fig. 1E) and soma (Fig. 1F). On day 2 (Fig. 1G-K), the NPC has divided asymmetrically, and the neuron’s soma is hidden by the NPC cell body in the z-projection, but the two closely apposed cell bodies that can be distinguished in the rotated X-projection (Fig1H), where the cell body of the neuron is more ventral. The axon of the newborn neuron uses the NPC as a scaffold and it projects along the radial process of the NPC for 37 µm before branching away and ending in small growth cones that contain mitochondria (asterisks in Figs. 1G,H). Both the NPC’s radial process and pial endfoot contain mitochondria (Fig. 1K). On day 3, the cell bodies of the NPC and its neuronal progeny have separated (Figs. 1L-O), and each extend a process for about 10 µm before they come together and the axon maintains its path along the radial process as on day 2, branching off at approximately the same position (arrow, Figs. 1L-N). Between day 2 and day 3, the axon has extended an additional 121 µm (it is outlined with the dotted line in Fig1N) and ends with a single growth cone. The NPC on day 3 maintains its pial endfoot on the dorsolateral tectal surface, and it maintains strong mitoEGFP expression throughout.

**Figure 1.**
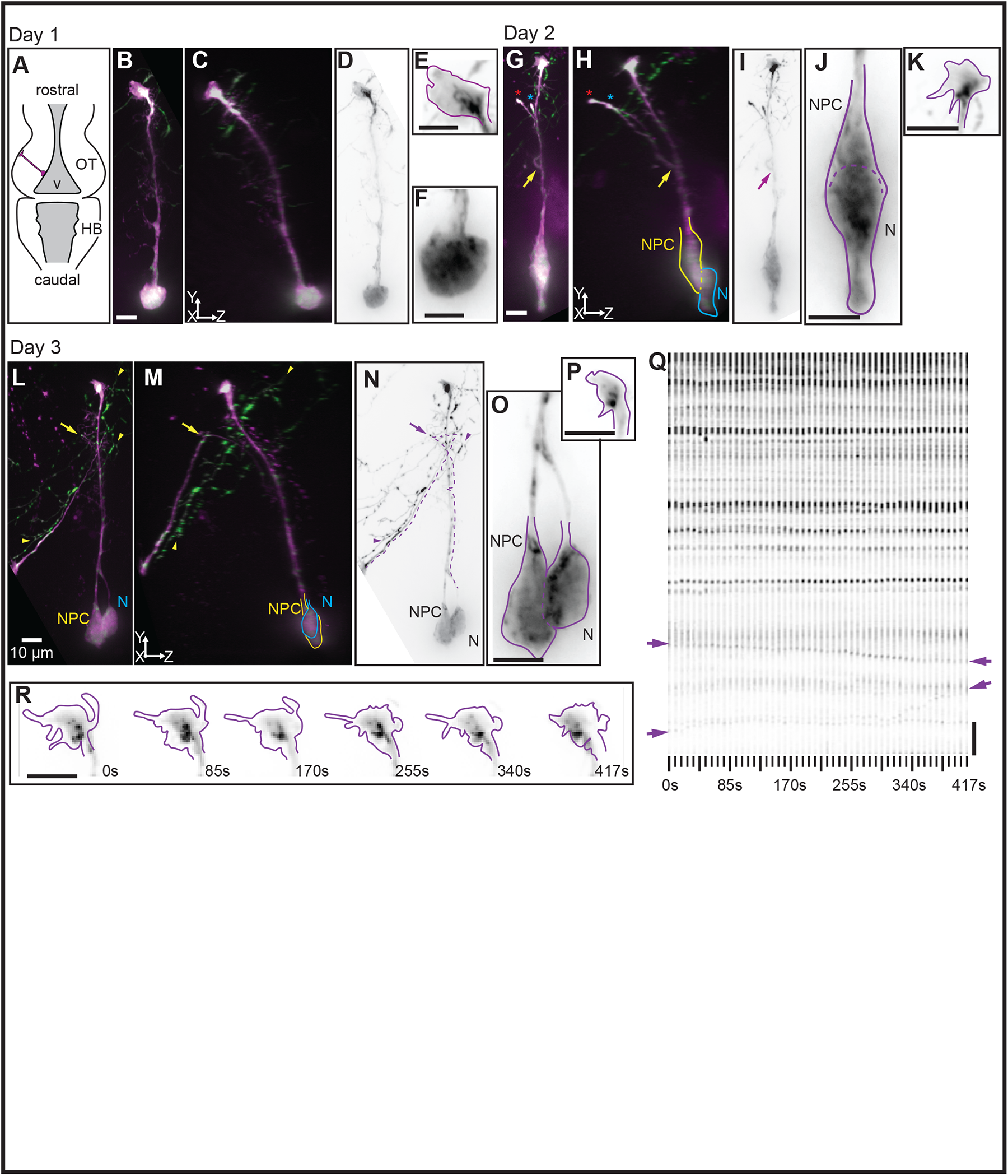
Tectal NPC expressing tagRFP-T (magenta) and mitoEGFP (green) imaged each day for 3 days. A) Position of tectal NPCs in the left lobe of the optic tectum in the brain OT: optic tectum; HB: hindbrain, v: ventricle. B-F) Tectal NPC on day 1. B) Maximum z-projection with the cell body of the NPC positioned toward the bottom of the image and the dorsolateral pial endfoot at the top. C) Same cell as in B, but the image is rotated 90° to produce the X-axis maximum projection revealing the dorsal curve of the NPC radial process. D-F) To better display the mitochondria, the mitoEGFP signal is grey-scaled and inverted in the z-projection of the entire cell (D), its outlined pial endfoot (E) and soma (F). G-K) The same NPC has divided 24 hours later. G is the maximum z-projection and H is the rotated X-oriented maximum projection with an outline defining the NPC and its neuron progeny. The arrow indicates the neuron’s axon branching away from the radial process of the NPC and the asterisks are the axon’s growth cone endings. I-K) Inverted mitoEGFP signal with views of the outlined somata of the NPC and the neuron (J) and pial endfoot (K). L-R) Day 3 image of the NPCs and neuron with z-projection of the cells in L, and x-rotated maximum projection in M. The arrow marks where the neuron’s axon has turned to project caudally along the dorsal surface of the tectum. N-P) Inverted mitoEGFP signal of the cell pair (N) and the dotted line is a slightly offset trace of the neuron’s axon. The two cell bodies are outlined (O) with mitochondria in the cell bodies and proximal processes of each cell. P) The pial endfoot of the NPC with mitochondria present. The arrowheads point toward an axon from a neuron elsewhere in the tectum that branches into this area. Q) The full 3D image was acquired in a rapid timelapse (50 images at 8.5 second intervals) and a kymograph of the mitoEGFP signal assembled to reveal the moving mitochondria (arrows) and the largely immobile mitochondria. In contrast, the mitochondria are very dynamic in the pial endfoot (Fig. R). Movies of the rotating cells and the rapid timelapse are available in the supplement. NPC: neural progenitor cell. N: neuron. Scale bars = 10 µm. 3D rotations of the cells on each day are available as Supplementary Videos (Figure1A-C and the timelapse on day 3).

To assess the level of fast dynamic movement of the mitochondria, the 3D confocal images were acquired every 8.5 sec for ∼7 minutes. The mitoEGFP signal along the radial process is arranged in the kymograph in Figure 1Q which reveals that most mitochondria are stationary and only 2 moved more than 5 µm during the timelapse (arrows, Fig. 1R). In contrast, the endfoot contains a dynamic network of mitochondria and over the course of the rapid timelapse the mitochondria move with every image captured (Fig1R). Videos 1-3 provide 3D rotated views of the NPC and its progeny on each day of the image series. Video 4 provides a rapid timelapse of the labeled cells on day 3 that shows the largely stable mitochondria in the NPCs but for a few small organelles that move along the radial process and the dynamic rearrangements that occur in the pial endfoot (Supplement Videos for Figure 1: day1, day2, day3, day3-timelapse).

### NPCs asymmetrically position mitochondria prior to cell division

Because the inheritance of mitochondria during cell division is associated with the fate of progeny (Aretz et al., 2020; Katajisto et al., 2015), we quantified the distribution of mitochondria in the NPCs prior to cell division to assess whether the mitochondrial topology signaled that the NPC was preparing to divide. We compared images of NPCs captured 24 hours before cell division (dNPC) and NPCs that remained quiescent during the 3 day timelapse (qNPC). Figures 2A-L show z-projections of a pair of NPCs where the NPC on the right divided on day 2 and the cell on the left remained quiescent. On each of the 3 days of the timelapse, both cells show the strong mitoEGFP expression in mitochondrial networks contained in their pial endfeet which are outlined and shown as inverted grey-scale images to reveal the fine details of the mitochondrial networks (Figures 2B, C, F, G, J, K). Fig2D, H and L are magnified images of the mitoEGFP signal within the cell bodies, revealing the stronger asymmetry in the distribution in the cell body of the dNPC one day before it divides. A 3D rotated video is provided in the supplement (Video Figure2_day1)

**Figure 2.**
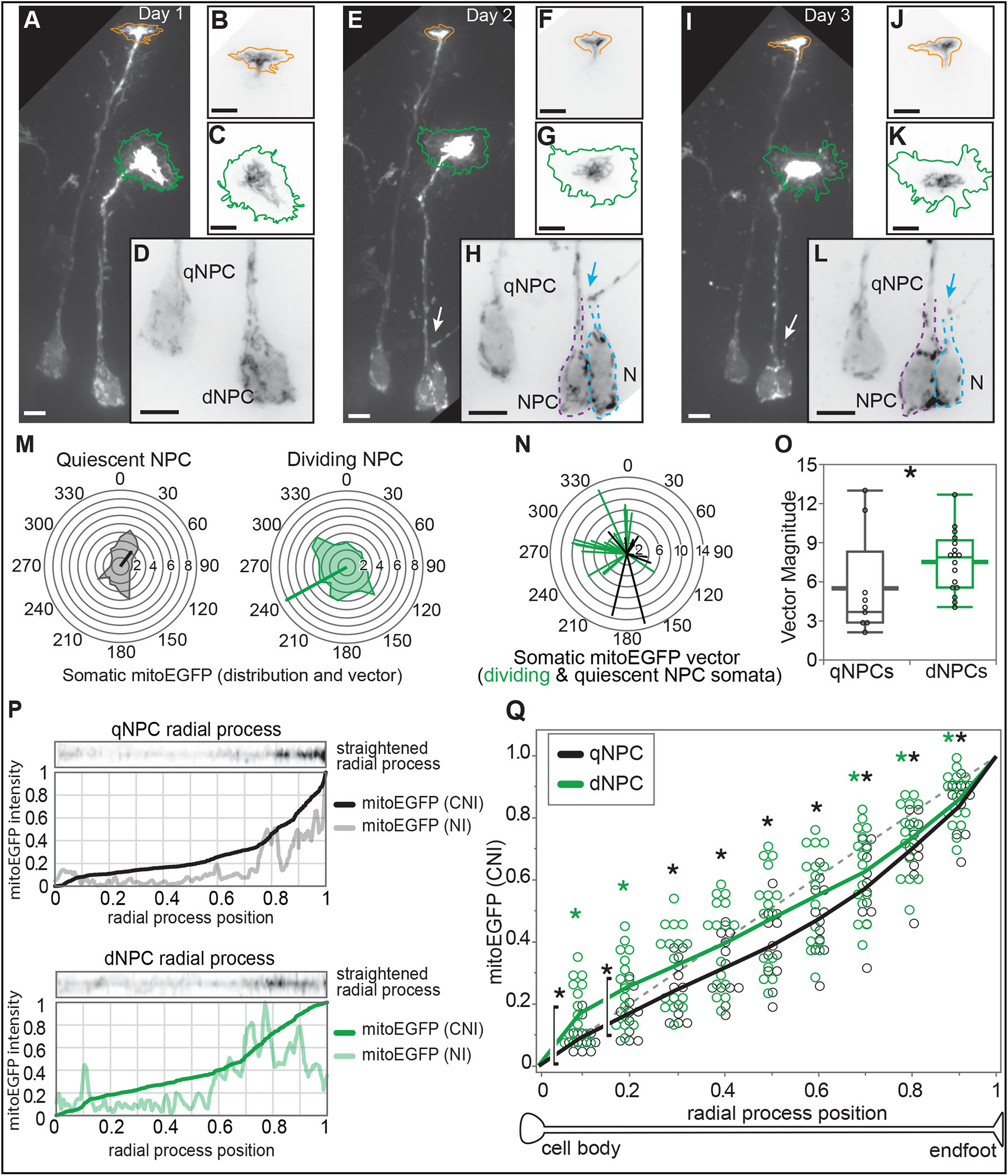
Mitochondria are unevenly distributed in tectal NPCs prior to cell division. A-L) A pair of mitoEGFP expressing NPCs in the right tectum imaged each day for 3 days. A, E, I) The maximum Z-projections of the 185 µm confocal stack of the cell pair are shown and the intensity is scaled so that the mitochondria are visible in the cell bodies. B,C, F, G, J, K) The pial endfeet of the qNPC (orange outline) and dNPC (green outline) shown in inverted grey scale to reveal the complex mitochondrial network in the endfeet. D, H, L) Inverted grey-scaled maximum projections through the cell bodies. The cell bodies of the parent NPC and its neuron progeny are outlined with dotted lines. On day 2 and 3, arrows point to the axon process of the neuron (N) branching away from the radial process of the NPC. Scale bars = 10 µm. M) Plots of the somatic mitoEGFP distribution are shown as shaded regions on the graph (grey, qNPC; green, dNPC) and the summed vector of the mitoEGFP signal for each cell is the line on each graph. N,O) The results of the somatic mitoEGFP distribution analysis revealing the magnitude and angle of the summed vector (N = 17 dividing and N = 9 quiescent NPCs). Comparing the vector magnitudes between the groups reveals a significantly greater bias in the distribution of somatic mitochondria for cells that will divide within 24 hours (p = Mann-Whitney U, p = 0.04). P) The maximum projections of the straightened radial processes for the qNPC and dNPC from day 1 are shown with the normalized mitoEGFP signal (grey and light green lines) and the cumulative normalized intensity of the mitoEGFP signal of the radial process (black and dark green lines). Q). The CNI values for all dNPCs (green) and qNPCs (black) with their mean values (green and black lines) at the different segments of the radial process reveals that NPCs that divide within 24 hours have greater levels of mitochondria positioned near the cell body. Diagram of an NPC with a radial process is stylized below the graph. The average distribution of mitochondria within the dNPCs’ radial processes was significantly different from the qNPCs (asterisk and brackets) and the asterisks above the data signify significant differences from the expected values if mitochondria were distributed equally (the dotted unitary slope line). Table 1 and 2 contain the statistical analysis of these data. 3D rotations of the cells day 1 are available as supplemental video of Figure 2.

To quantify the mitochondrial distribution in the cell bodies of dividing and quiescent NPCs, we developed a morphometric analysis method to calculate the magnitude and orientation of the mitoEGFP signal. This method allowed us to quantitatively assess the distribution of the mitoEGFP reporter and make comparisons between cell groups. Fig2M displays the spatial distribution of the mitoEGFP signal (the shaded area of the figures) and the line is its summed vector and angle (where 0° is the position where the radial process emerges from the cell body). Because there is no strong aggregation of mitochondria in the soma of the qNPC, its vector value is 2.1 at 37°. In comparison, the mitochondria in the cell body of the dNPC were aggregated which produced greater vector magnitude of 8.1 at 241° (Fig. 2M). In total, we measured the somatic mitoEGFP distribution for 18 dNPCs (green) and 9 qNPCs (black; Figure 2N). The mean vector value for the dNPC group, 7.5 ±0.57 (N =17), was more than 30% greater and significantly different from the 5.5±1.32 (N =9) mean for the qNPC cells (Mann-Whitney U test, p = 0.04). These data show that NPCs that are preparing to divide have a biased aggregation of mitochondria in their somata compared to the quiescent cells.

The radial process of NPCs is the cellular link between the relatively large reservoir of mitochondria in the pial endfoot (Figs2A-K) and their cell bodies. We measured the mitoEGFP distribution in the radial process in order assess whether mitochondria were repositioned toward the cell body, increasing the likelihood they could be inherited by the NPC’s progeny during cytokinesis. Fig2P shows the straightened 2D maximum projection of the radial processes of the two NPCs (inverted greyscale images). The mitoEGFP fluorescence was measured from the cell body to the pial endfoot from the 3D confocal image stack and the values normalized to the maximum. In order to make comparisons between cells, these values were plotted against the normalized radial process length where 0 is the point near the cell body and 1 is the distal end of the radial process near the pial endfoot (Figure 2P, light grey and light green lines of the normalized intensity; NI). Next, we summed the mitoEGFP intensity to derive the cumulative normalized intensity (CNI) scaling it from 0 to 1 (black and green lines). In short, the CNI metric reports the normalized distribution of mitochondria for each cell so that comparisons between the quiescent cells and those that divided within 24 hours could be made.

For both the dNPC and the qNPC shown in Figure 2, mitochondria are not equally distributed along their radial processes. In the qNPC, 50% of the mitoEGFP signal is contained within the distal 18% of the radial process (near the pial endfoot). In comparison, the mitoEGFP signal in the radial process of the dNPC was more evenly distributed, with 50% of the mitoEGFP signal detected in the proximal 69% of the radial process (Fig.2P). Figure 2Q provides the mean CNI for the qNPCs (black line; n = 12) and dNPCs (green line; N =16) at each of the segments of the radial process along with CNI values from individual cells (green, dNPCs and black, qNPCs). Comparing the dNPC CNI values against the expected values if mitochondria were evenly distributed (the grey dotted unitary line, Fig.2Q) revealed that dNPCs had significantly greater than expected levels of mitochondria at nearest the cell body (Wilcoxon signed rank test, p = 0.01 and p = 0.05 at positions 0.1 and 0.2; green asterisks, Figure 2Q) and deviated from the expected values at the more distal positions of 0.7 and 0.8 (Wilcoxon signed rank test, p = 0.04 and p = 0.03; see Table 1). In contrast, the average qNPC CNI was evenly distributed for the first two segments of the radial process but was less than expected for the remaining distal 80% of the radial process (black asterisks, Wilcoxon signed rank tests, p = 0.002 to 0.05, see Table 1). It was also telling that the dNPC CNI values were also significantly different from the qNPC values for the most proximal 20% of the radial process (asterisk with brackets, Figure 2Q; Mann Whitney U test, p = 0.01 and p = 0.04). Together, the CNI analysis reveals that when NPCs are within 24 hours of dividing, mitochondria are repositioned closer to the cell body, suggesting that they are moved so that they can be inherited by the cell progeny.

**Table 1.** Table of statistical measurements used in the somatic mitoEGFP distribution analysis.

| Cell type | N | Mean | Median | S.D. | Lower 95% CI | Upper 95% CI | Mann-Whitney U |
| --- | --- | --- | --- | --- | --- | --- | --- |
| Dividing NPC | 17 | 7.89 | 8.0 | 2.70 | 6.54 | 9.23 | **p = 0.0328** |
| Quiescent NPC | 9 | 5.50 | 3.7 | 3.96 | 2.46 | 8.54 |  |

The Mann-Whitney U values in red indicate that there was a significant difference in the distribution of mitoEGFP in the cell bodies of dNPC and the qNPC groups.

**Table 2.**
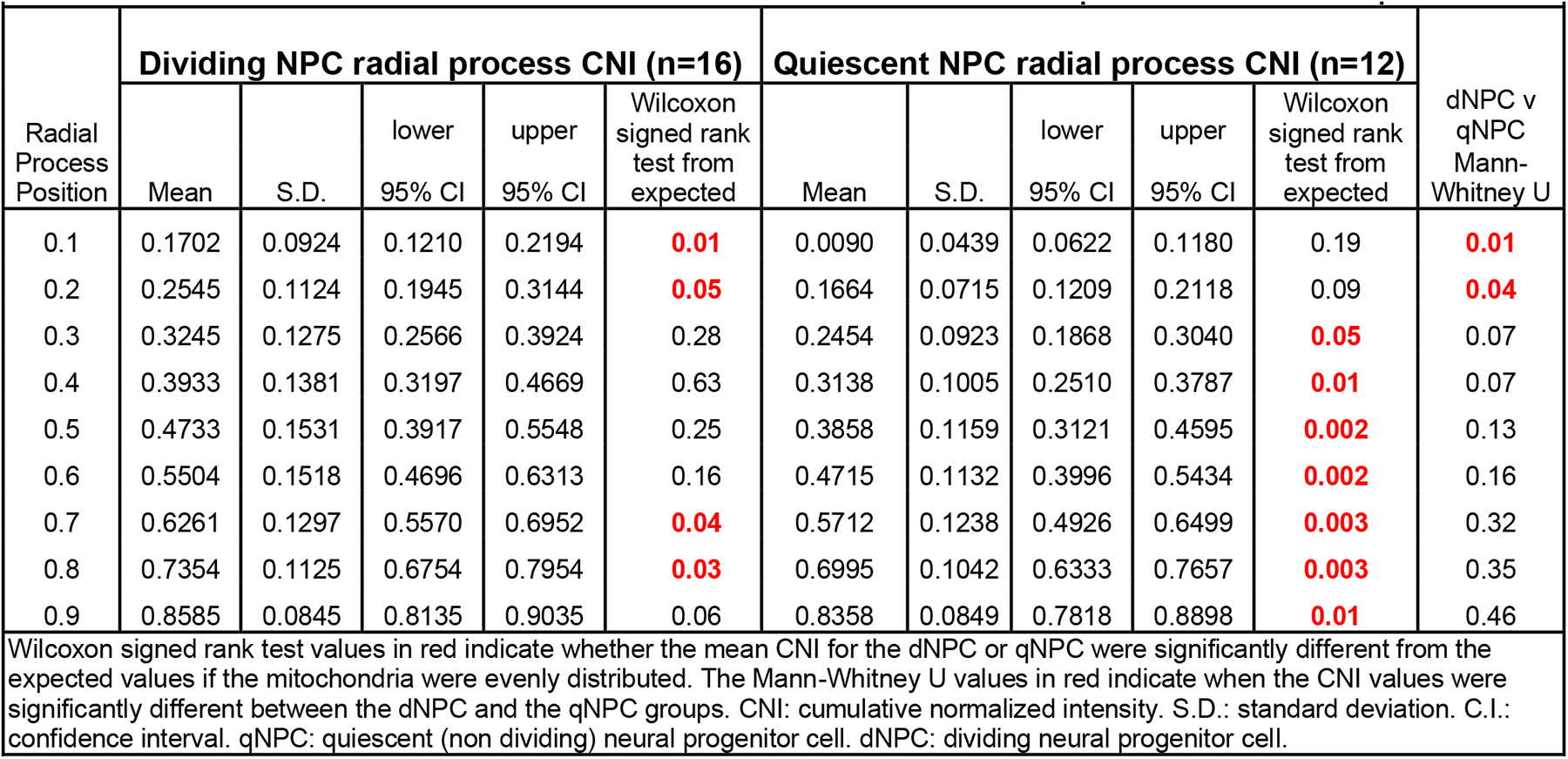
Table of statistical measurements used in the radial process CNI comparisons.

| Radial Process Position | Dividing NPC radial process CNI (n=16) |  |  |  |  | Quiescent NPC radial process CNI (n=12) |  |  |  |  | dNPC v qNPC Mann-Whitney U |
| --- | --- | --- | --- | --- | --- | --- | --- | --- | --- | --- | --- |
|  | Mean | S.D. | lower 95% CI | upper 95% CI | Wilcoxon signed rank test from expected | Mean | S.D. | lower 95% CI | upper 95% CI | Wilcoxon signed rank test from expected |  |
| 0.1 | 0.1702 | 0.0924 | 0.1210 | 0.2194 | **0.01** | 0.0090 | 0.0439 | 0.0622 | 0.1180 | 0.19 | **0.01** |
| 0.2 | 0.2545 | 0.1124 | 0.1945 | 0.3144 | **0.05** | 0.1664 | 0.0715 | 0.1209 | 0.2118 | 0.09 | **0.04** |
| 0.3 | 0.3245 | 0.1275 | 0.2566 | 0.3924 | 0.28 | 0.2454 | 0.0923 | 0.1868 | 0.3040 | **0.05** | 0.07 |
| 0.4 | 0.3933 | 0.1381 | 0.3197 | 0.4669 | 0.63 | 0.3138 | 0.1005 | 0.2510 | 0.3787 | **0.01** | 0.07 |
| 0.5 | 0.4733 | 0.1531 | 0.3917 | 0.5548 | 0.25 | 0.3858 | 0.1159 | 0.3121 | 0.4595 | **0.002** | 0.13 |
| 0.6 | 0.5504 | 0.1518 | 0.4696 | 0.6313 | 0.16 | 0.4715 | 0.1132 | 0.3996 | 0.5434 | **0.002** | 0.16 |
| 0.7 | 0.6261 | 0.1297 | 0.5570 | 0.6952 | **0.04** | 0.5712 | 0.1238 | 0.4926 | 0.6499 | **0.003** | 0.32 |
| 0.8 | 0.7354 | 0.1125 | 0.6754 | 0.7954 | **0.03** | 0.6995 | 0.1042 | 0.6333 | 0.7657 | **0.003** | 0.35 |
| 0.9 | 0.8585 | 0.0845 | 0.8135 | 0.9035 | 0.06 | 0.8358 | 0.0849 | 0.7818 | 0.8898 | **0.01** | 0.46 |

Wilcoxon signed rank test values in red indicate whether the mean CNI for the dNPC or qNPC were significantly different from the expected values if the mitochondria were evenly distributed. The Mann-Whitney U values in red indicate when the CNI values were significantly different between the dNPC and the qNPC groups. CNI: cumulative normalized intensity. S.D.: standard deviation. C.I.: confidence interval. qNPC: quiescent (non dividing) neural progenitor cell. dNPC: dividing neural progenitor cell.

### Overexpression of mitochondrial biogenesis regulator, pgc1a induces neurogenic cell division

These morphometric analyses revealed that NPCs within 24 hours of dividing have unequal distribution of mitochondria within their cell bodies and radial processes. It is consistent with other studies that show that the inheritance of mitochondria regulate how the NPCs divide and the fate of their progeny (Dohla et al., 2022). We were interested in testing whether increased levels of mitochondria could alter the cell division fate of the NPCs by overexpressing the “master regulator” transcription factor of mitochondrial biogenesis, peroxisome proliferator-activated receptor gamma coactivator 1-alpha (ppargc1a, which encodes PGC-1a). PGC-1a controls the expression of transcription factor networks that regulate mitochondrial biogenesis and oxidative metabolism (Scarpulla et al., 2012) and it is highly expressed during neuron differentiation (Zheng et al., 2016) where it regulates mitochondrial genes and mitochondrial respiration levels (Cowell et al., 2009) (Rosenkranz et al., 2021; Wareski et al., 2009). We electroporated tectal NPCs with expression vectors containing ppargc1a and cytosolic tagRFP-T in order to prematurely induce mitochondrial biogenesis in the NPCs and test whether mitochondria could alter the fate of the NPCs.

An NPC overexpressing ppargc1a is shown in Figure 3. Over the course of 3 days this cell divides symmetrically to produce an additional NPC before dividing asymmetrically to generate a new neuron. On day 1, the NPC’s 148 µm radial process terminates in a single 10 µm diameter endfoot on the ventrolateral surface of the brain (arrows, Fig. 3A-F). Figure 2D shows the mito-EGFP signal alone (inverted grey scale), and like control cells (Figs 1,2) mitochondria are found throughout all compartments of the cell including the expansive pial endfoot (Fig. 3E) and cell body (Fig.3F). In this cell, the distal third of the radial process contains the strongest mitoEGFP expression, compared to the lower and aggregated mitochondrial signal in the cell body which is more easily seen in the magnified views (Fig. 3F). A mitochondrial network is visible in the grey-scaled images of the pial endfoot (Fig. 3E), highly characteristic of the tectal NPCs. Tectal NPCs often have dynamic filopodial branches along the radial process (Sild et al., 2016; Tremblay et al., 2009). Fig. 3B and C provide rotated projections to reveal the radial process, the many filopodia and the distal pial endfoot. Most filopodia have low expression of mitoEGFP, but the asterisks label a mitoEGFP-expressing branch with distinct mitochondria that are persistent throughout the 3 days (asterisks, Fig. 3).

**Figure 3.**
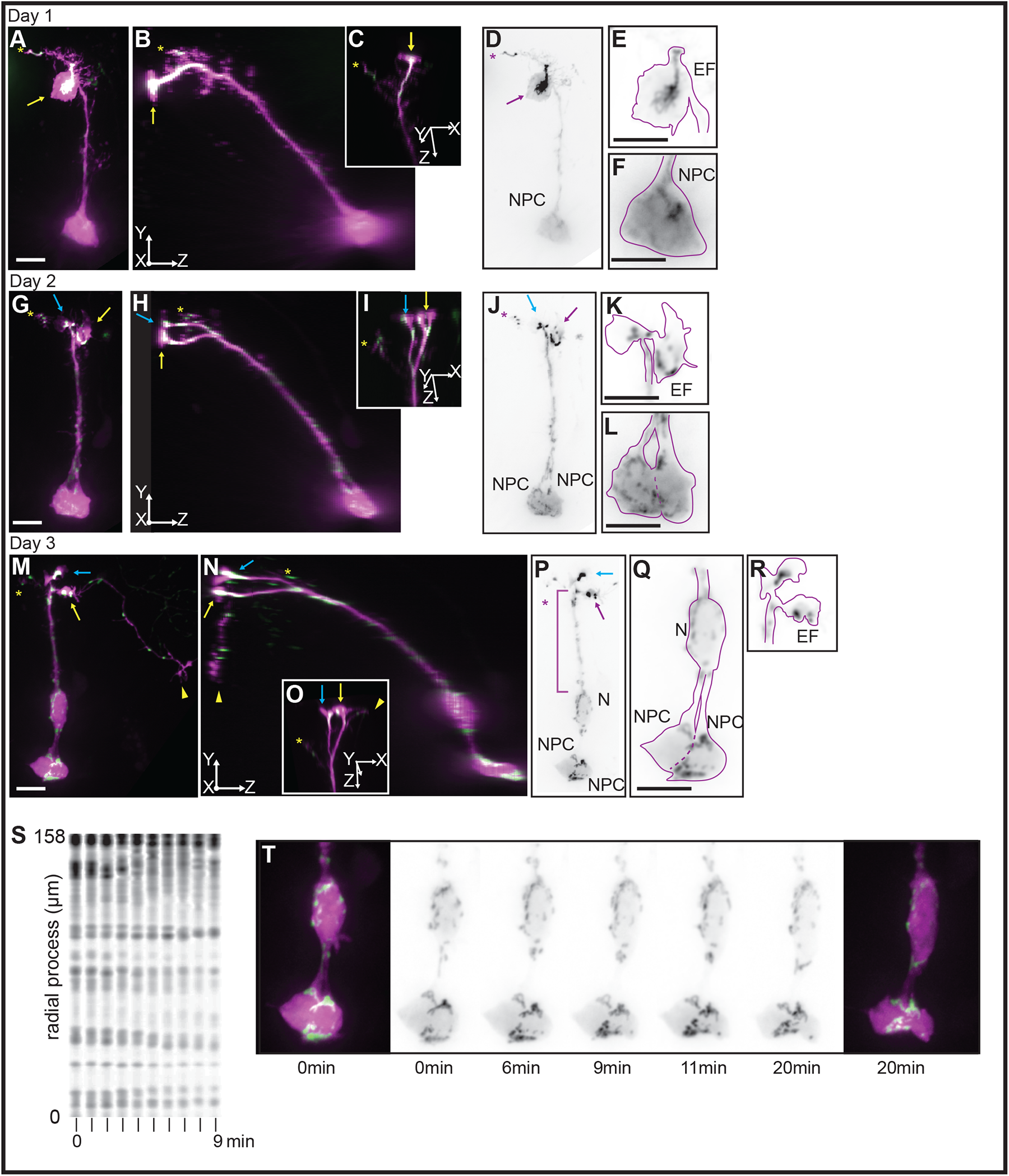
Proliferating tectal NPC overexpressing ppargc1a, tagRFP-T (magenta) and mitoEGFP (green) imaged each day for 3 days. A, G, M) Z-projections of the NPC and its progeny on day 1-day 3. B, H, N) X-rotated projection of the cells with the dorsolateral endfoot on the left edge. C, I, O) Rotated maximum projections of the pial endfeet are shown in C, I and O). On day 1, the large endfoot (yellow arrow) contains a mitochondrial network (E). There are fine filopodial branches near the distal end of the NPC, and the longest branch contains mitochondria (asterisk). This branch persists over the 3 days. The cell body has relatively few mitochondria on day 1(F), but on day 2, the cell has divided (L) and the cell bodies contain more mitochondria. The radial process of the second cell has grown along the original NPC until it reaches the dorsolateral surface where it branches and makes its own endfoot (teal arrow). There is a second endfoot (teal arrows, G-J). On day 3, there is a newborn neuron, and its cell body is ∼15 microns along the radial processes of the two NPCs. Q) the outlined cell bodies of the 3 cells revealing the differences in the number of mitochondria between the cells. The pial endfeet of the 2 NPCs contain mitochondria, and the axon of the neuron has grown along the radial processes to reach the dorsal surface of the tectum before it has branched and grown along the surface of the tectum (arrowhead, M, N). 3D rotations of the cells on day 3 are available as Supplemental video for Figure 3. Scale bars = 10 µm.

By day 2, the NPC has divided to produce two NPC cell bodies (outlined in Fig. 3L). Each cell contributes a radial process that emerges from the cell bodies before they fasciculate, extend together toward the ventrolateral surface and split apart to interface with the surface of the tectum. The new NPC’s pial endfoot has grown (teal arrow, Fig. 3G-I) and the original NPC’s pial endfoot that was visible on day 1 has reduced in size and split so that it terminates in two smaller endfeet (yellow arrow, Fig.3 G-I). The morphological changes to the pial endfeet are most obvious in the rotated magnification in Fig.3I. The mitochondrial network in these distal endfeet has simplified by day 2 and individual organelles are visible in the outlines of the endfeet (Fig. 3K). The distal bias in mitochondria has decreased and the mitochondria are more equally distributed throughout the cells on day 2.

On day 3 a newborn neuron appears, and its cell body is ∼15 µm away from the ventricle where the cell bodies of the two NPCs from day 2 remain (Fig3M, N, Q). The axon of this neuron and radial processes of the NPCs are tightly fasciculated and cannot be distinguished until they reach the ventrolateral surface of the tectum where the two NPCs separate and each terminate with their endfeet (teal and yellow arrows, Fig. 3M-O). The axon branches off from the NPC endfoot marked with the yellow arrow (Figs. 3M, N) and projects caudally along the surface of the tectum. The axon contains mitochondria and ends with a branching growth cone (arrowhead, Figs. 3M-O). Mitochondria persist in the NPC endfeet (Fig.3R) and are found in the somata of all 3 cells (Fig3 Q and T). Interestingly, on day 3, compared to the soma of the right NPC, the cell on the left (Fig3. M, P, Q) has fewer mitochondria that are accumulated near where the radial process emerges from the cell body. Similar to the control cells (Fig. 1Q), the kymograph of the radial process (length indicated with the bracket, Figure P) revealed that after a 9 minute timelapse the mitochondria in the radial processes are largely stationary (Fig. 3S). The image series of the cell bodies (Fig. 3T) shows that the mitochondria occupy largely the same positions over the 20 minute period. The mitochondria and fine details of this cell group can be more easily inspected in Supplement Video (Figure3_day3.mp4) which provides views of the rotating cell group on day 3. These data revealed that distribution and number of mitochondria in the NPCs overexpressing ppargc1a was not notably different from the control NPCs.

Figure 4 summarizes the lineages and division events of 30 control NPCs and 30 NPCs that overexpress ppargc1a. The control group expressed cytosolic RFP alone or in combination with mito-EGFP, and the ppargc1a NPCs overexpressed ppargc1a in addition to the fluorescent proteins. For both NPC groups, the NPCs that remained quiescent and did not divide during the 3 day window were the smallest portion of the cell lineages; they made up 23% (7/30) of the control NPCs and 20% (6/30) of ppargc1a NPC lineages. Overexpression of ppargc1a increased the proportion of cells that divided asymmetrically and generated neurons during the 3 day window. We found that compared to the 43% (13/30) of control NPC lineages that produced neurons, 63% (19/30) of ppargc1a-expressing NPCs divided asymmetrically. With this increase in neurogenesis induced by ppargc1a expression, only 17% (5/30) NPCs remained in the symmetrically dividing progenitor pool compared to the 30% (10/30) of control NPCs. These changes in the dividing NPC lineages were significantly different between the two cell groups (Pearson Chi-Square, p = 0.03). The consequence of overexpressing ppargc1a was to significantly increase asymmetric cell division and increase the number of neurons born over 3 days compared to the control cells. On day 3, 30% (16/53) of the cells generated by the 30 control NPCs were neurons compared to the 44% (27/61) neurons that were generated by the 30 NPCs expressing ppargc1a (Pearson Chi-Square, p = 0.02). In short, increasing levels of mitochondrial biogenesis through the overexpression of ppargc1a induced neurogenesis in the tectal NPCs.

**Figure 4.**
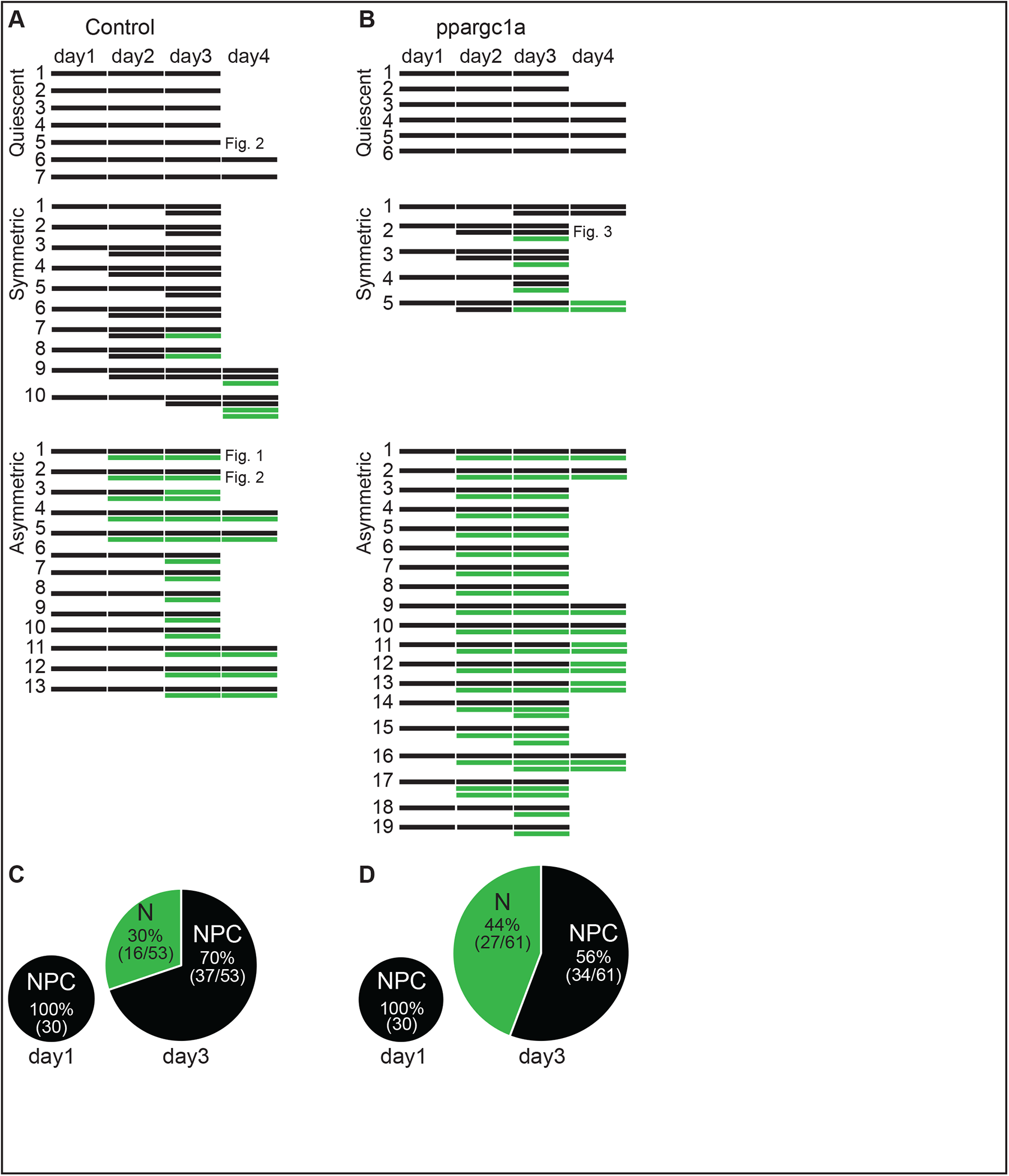
Neurogenic cell proliferation decreases in NPCs treated with ppargc1a morpholinos. A-D) Maximum z-projections of the left tectal lobe imaged 1 day and 3 days after co-electroporation of pSox2-bd::EGFP and the control morpholino (A,B) or the ppargc1a morpholino, which blocks translation (C,D). The asterisks indicate pial endfeet of NPCs which are less abundant in the control-MO group on day 3 as the number of branching neurons increases. The arrow points to a cluster pial endfeet from of ppargc1a-treated NPCs that have multiplied by day 3. The tectum is outlined with the dotted line and the scale bar = 20 µm. E-H) Summary graphs of the changes in the percentages of neurons and NPCs for the control (E,F) and ppargc1a-MO (G,H) treated groups. The asterisks indicate a significant change between day 1 and day 3 (Wilcoxon Signed Rank test, p < 0.05). I-K) By day 3, NPCs treated with the ppargc1a-MO generated significantly fewer neurons than the controls on by day 3 (I) and more NPCs (J) than the control MO group by day 3 (p<0.05; Mann-Whitney U test). K) Compared to NPCs treated with the control MO, ppargc1a MO treated NPCs divided less over the 3 days and generated fewer cells overall (p<0.05; Mann-Whitney U test). Graphs show the individual data points, the mean (thick horizontal line) and quantile boxplots. Asterisk indicates significant differences. The mean values, standard errors, 95% confidence intervals and p-values are reported in Table 3.

### Inhibiting PGC-1a expression limits neurogenesis and cell proliferation

We next assessed whether interfering with mitochondrial biogenesis by knocking down PGC-1a would alter expected levels of cell proliferation and neurogenesis. We used an antisense morpholino oligonucleotide (MO) designed to interfere with translation initiation of ppargc1a and a control MO that does not target any known *Xenopus* sequence. When administered using electroporation, morpholinos are effective at interfering with protein expression without the potential off-target effects that can occur when high concentrations of morpholinos are used to create whole-animal knockdown (Bestman and Cline, 2020). After co-electroporating pSox2-bd::eGFP along with lissamine-tagged control MO or ppargc1a MO, we acquired full 3D confocal stacks of all labeled cells in the tectum. On day 3, the same image was acquired and the percent changes in the total number of cells and in the proportions of NPCs and neurons were measured. Examples of EGFP-positive cells from the control MO (Figs. 5A-B) and ppargc1a MO (Figs. 5C-D) groups are shown as maximum z-projections of the cells in left tectal lobes. The asterisks indicate individual pial endfeet of NPCs and the arrow points toward a group of pial endfeet. Figures 5E-H show the percent change in NPCs and neurons in 11 control-MO and 20 ppargc1a-MO tectal lobes on day 1 and day 3. On day 1, there are no significant differences in the percentages of neurons or NPCs for the two groups (Table 3). Comparing the percentages of NPCs and neurons on day 1 and day 3 showed that both the control and ppargc1a-MO groups yielded decrease in NPCs (Figs. 5F, G; Table 3) and a corresponding a significant increase in neurons (Figs. 5E, G; Table 3) as the cells differentiated. Knocking down ppargc1a interfered with the level of neurogenesis (Fig5. I); while the percentage of neurons in the control group increased to 71.1±5.1% on day 3 (with a corresponding in the percentage of NPCs to 28.2±5.1%), the ppargc1a MO group did not match these levels. The ppargc1a MO tecta yielded 54.0±5.2% neurons and 46.0±5.2% NPCs on day 3, both were significantly different from the control levels (Figs. 5 I, J; Mann Whitney U test, p = 0.05; Table 3). Knocking down PGC-1a expression also significantly reduced the total number of cells generated over the 3 day window (Fig. 5K). Where the control group yielded an 82.3±20.3% increase in the number of cells generated, the ppargc1a-MO expressing group had significantly fewer cells generated, yielding just a 30.2±7.7% increase (Mann-Whitney U; p = 0.02; Table 3). Together these results show that interfering with mitochondrial biogenesis by knocking down PGC-1a significantly increased the number of NPCs in the tectum and this resulted in the production of few neurons.

**Figure 5.**
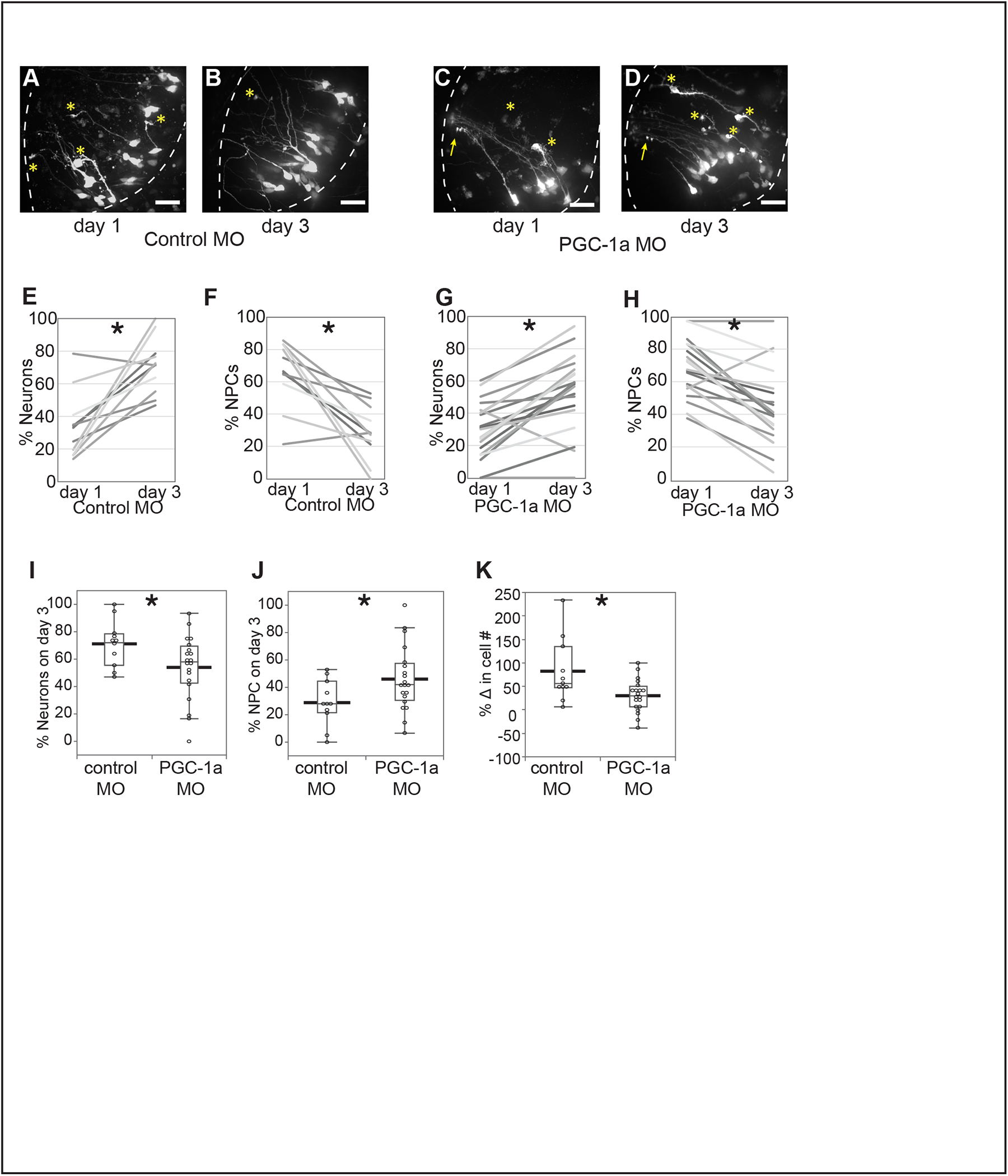
Neurogenetic cell proliferation increases with ppargc1a overexpression. A-B) Lineages of 3D confocal images acquired each day for 3 or 4 days of 30 control NPCs expressing only fluorescent proteins (A) and 30 NPCs also overexpressing ppargc1a (B). Each lineage begins with a single NPC (black bar) and the addition of a second bar marks that the NPC divided either symmetrically (the addition of a black bar) or asymmetrically when a neuron is generated (the addition of a green bar). The cells shown in Figures 1-3 are indicated. NPCs that overexpress ppargc1a are more likely to divide asymmetrically compared to the control group (Pearson Chi-square p = 0.02). C-D) Proportions of the cells on day 1 and day 3 for the control (C) and ppargrc1a-expressing NPCs (D). The size of the pie chart indicates the relative change in total cell numbers. C) The control lineages yielded a 76% increase in cells on day 3 with 30% (16/53) neurons and 70% (37/53) NPCs. D) Compared to the control group, NPCs overexpressing ppargc1a doubled the number of cells generated over 3 days. They produced 44% (27/61) neurons and 56% (34.61) NPCs by day 3, yielding significantly greater levels of neurogenesis than the control group (Pearson Chi-square p = 0.02).

**Table 3.** Table of statistical measurements used in the analysis of cell proliferation in control and PGC-1a morpholinos in tectal NPCs.

| Experiment | N | % Neurons day 1 |  |  |  |  | % Neurons day 3 |  |  |  |  | Wilcoxon Signed Rank p-value |  |
| --- | --- | --- | --- | --- | --- | --- | --- | --- | --- | --- | --- | --- | --- |
|  |  | Mean % | S.D. | Lower 95% CI | Upper 95% CI | Mann Whitney U | Mean % | S.D. | Lower 95% CI | Upper 95% CI | Mann Whitney U |  |  |
| Control MO | 11 | 34.2 | 20.1 | 20.7 | 47.8 | p=0.88 | 71.2 | 16.8 | 59.9 | 82.4 | p = 0.05 | p=0.0117 |  |
| PGC-1a MO | 20 | 31.4 | 16.1 | 23.6 | 39.1 |  | 54.0 | 23.5 | 43.0 | 65.0 |  | p<0.0001 |  |
| Experiment | N | % NPCs day 1 |  |  |  |  | % NPCs day 3 |  |  |  |  | Wilcoxon p- value |  |
|  |  | Mean % | S.D. | Lower 95% CI | Upper 95% CI | Mann Whitney U | Mean % | S.D. | Lower 95% CI | Upper 95% CI | Mann Whitney U |  |  |
| Control MO | 11 | 65.8 | 20.1 | 52.2 | 79.3 | p=0.71 | 28.8 | 16.8 | 17.6 | 40.1 | p = 0.05 | p=0.002 |  |
| PGC-1a MO | 20 | 70.2 | 17.2 | 62.2 | 78.2 |  | 46.0 | 23.5 | 35.0 | 57.0 |  | p<0.0001 |  |
| Experiment | Day 1 total cells |  |  |  | Day 3 total cells |  |  |  | Percent Change in Cell # |  |  |  | Mann Whitney U |
|  | Mean Cell # | S.D. | Lower 95% CI | Upper 95% CI | Mean Cell # | S.D. | Lower 95% CI | Upper 95% CI | % Cell # Change | S.D. | Lower 95% CI | Upper 95% CI |  |
| Control MO | 14.7 | 8.3 | 9.1 | 20.3 | 24.9 | 12.8 | 16.3 | 33.5 | 82.3 | 67.2 | 37.2 | 127.5 | P=0.02 |
| PGC-1a MO | 18.9 | 7.9 | 15.2 | 22.6 | 23.9 | 10.4 | 19.1 | 28.7 | 30.2 | 34.5 | 14.1 | 46.3 |  |

The Mann-Whitney U values in red indicate that there was a significant difference in the percent of NPCs or neurons between the control and ppargc1a MO groups. Wilcoxon values in red indicate significant difference in the percent of neurons or NPCs between day 1 and day 3. MO: morpholino. S.D.: standard deviation. C.I.: confidence interval.

## DISCUSSION

The nervous system demands more from mitochondria than any other tissue (Engl and Attwell, 2015) and therefore its development is also the most susceptible and sensitive to mitochondrial dysfunction (Koenig, 2008). A critically important step in healthy brain development is the inheritance of mitochondria through cell division; the mitochondria that newborn neurons inherit will be the only source of the organelle that must serve that neuron for decades. While it is clear that mitochondrial dysfunction alters neurogenesis (Arrazola et al., 2019; Xavier et al., 2016), at present we lack a full understanding how mitochondria are regulated by the NPCs or inherited by their progeny. Much of what is currently known about the role that mitochondria play in cell proliferation comes from measurements of cell populations or from cultured cells in vitro. To our knowledge, our study is the first to take a longitudinal imaging approach with single cell resolution that investigates mitochondria throughout the full NPC architecture prior to and after they divide. We demonstrate that mitochondria are not equally distributed in the radial process and our image analysis results reveal a distinct bias in the cell bodies of NPCs as they prepare to divide. By targeting individual NPCs with tools to increase and decrease levels of PGC-1a we found that when mitochondrial biogenesis is induced in NPCs, they are significantly more likely to produce neurons and that by suppressing PGC-1a, both the likelihood of NPC proliferation and neurogenesis decrease. Together our data show that mitochondria contribute to the regulation of NPC cell division and fate of their progeny.

### Mitochondria distribution and movements in tectal NPCs

We find that tectal NPCs, like cortical NPCs have abundant mitochondria that distribute throughout the cells (Figs 1-3), which is striking given that proliferating cells do not to generate ATP (Agathocleous et al., 2012; Petridi et al., 2022). The elongated, polarized NPCs are unique brain cells in that their cell bodies or apical endfoot physically interfaces with inner ventricle of the brain, but they also contact the outer surface though interactions of their pial endfeet with vasculature and the developing meninges. Compared to the other compartments of the NPCs, we find the most complex mitochondrial networks are found in the pial endfeet (Fig. 1E, Fig. 2B, C, Figs. 3D-E). Though at the distal extreme from the cell body of a NPC, the pial endfoot of NPCs is responsible for a range of processes. Cell-cell signaling that induces cell division is mediated by the pial endfeet of NPCs (Pilaz and Silver, 2017) and they are sites of active endocytosis (Lau et al., 2017) and local protein synthesis for the NPCs (Pilaz et al., 2016). NPC endfeet direct the growing retinal ganglion cell axons as they enter the tectum (Vanselow et al., 1989), and the distal end of the NPCs have highly dynamic and motile filopodia that are responsive to changes in the visual experience of the animals (Sild et al., 2016; Tremblay et al., 2009). It is in this cellular compartment of the NPCs where we find the highest density of mitochondria (Figs1-3), similar to what Rash et al., (2018) describe in the NPCs of the developing mouse cortex. Unlike our results, where mitochondria in the radial process are largely stationary (Figs 1Q and 3S), Rash et al. report that ∼90% of the mitochondria in the radial processes of mouse cortical NPCs are motile. Furthermore, they find that mitochondrial movements are inversely related to calcium transients in the cells (Rash et al., 2016; Rash et al., 2018). Calcium transients occur less frequently in the tectal NPCs (Tremblay et al., 2009) compared to those recorded from NPCs in rodent cortex (Rash et al.; Weissman et al., 2004), which may explain the relative immobility of the mitochondria in this study. Culture conditions of cortical slice culture require elevated glucose levels compared to the in vivo environment (Silver and Erecinska, 1994), which may also contribute to the differences we observe. The relatively low levels of directed mitochondrial movement we find are more in line with the relatively small portion (10-40%) that move in axons when measured *in vivo* compared to the higher proportions that move *in vitro* (Misgeld and Schwarz, 2017). The most accurate estimations of mitochondria movement would be made by labeling the organelles with photoactivatable and photoconvertible fluorescent reporters (Matsuda and Nagai, 2014) so that individual mitochondrion can be tracked as they move, fuse with others and undergo fission, similar to the experiments now being done to measure the transport of mitochondria in axons (Wehnekamp et al., 2019) or track the differential inheritance of mitochondria of dividing cells in culture (Katajisto et al., 2015).

In order to compare NPCs that were imaged in the day preceding cell division versus those that remained mitotically quiescent, we use two morphometric analysis approaches to describe the mitochondrial topology in the cells. Examination of the distributions of mitochondria in the cell body revealed that the majority of the quiescent NPCs showed no bias in the distribution of their mitochondria, compared to the cells preparing to divide. Prior to cell division, NPCs accumulate mitochondria away from the ventricular and toward the radial process (Fig. 2) and this results in a significant bias in the mitochondrial distribution. Furthermore, comparisons of the cumulative normalized intensity of the mitochondria within the radial process also detected a significant bias in mitochondrial distributions. NPCs appear to prepare for cell division by increasing the relative levels of mitochondria near the cell body (Figs. 2P-Q). Further analysis is needed to understand whether the unequal distributions of mitochondria are the result of new mitochondrial biogenesis from existing mitochondria located in and around the NPC cell bodies, if they result from the directed movement of mitochondria to those positions in the cell, or a combination of each type of regulation.

In light of these results, it is interesting to consider the recent work showing that mitochondria are partitioned so that they are not only unequally inherited by the progeny of dividing stem cells *in vitro*, and functionally distinct mitochondria are inherited by the cells destined to differentiate (Dohla et al., 2022; Katajisto et al., 2015). Stem cells that inherit younger mitochondria are more likely to remain in the stem cell pool compared to sibling cells that inherit the older mitochondria, which are more likely to differentiate (Dohla et al., 2022; Katajisto et al., 2015). Older mitochondria are more likely to have a higher mutation levels due to damaging reactive oxygen species that are generated with ATP production, and by biasing their inheritance to the differentiated cells, the dividing stem cells are spared these potentially damaged organelles (Dohla et al., 2022; Katajisto et al., 2015). Older mitochondria may be less efficient in producing ATP (they have a lower membrane potential), but this inefficiency in ATP production may free up substrates normally used in mitochondrial oxidative phosphorylation for other biochemical pathways that could be critical to differentiating and developing neurons (Spinelli and Zaganjor, 2022; Vander Heiden et al., 2009). Given that millions of neurons are generated each hour during embryonic neurogenesis (Silbereis et al., 2016), mechanisms that could direct the inheritance of damaged mitochondria to the differentiated neuron progeny would prevent the further propagation of potential mutation as the NPCs continue to divide.

### PGC-1a-mediated changes to mitochondria in NPCs directs symmetric or asymmetric cell divisions

The prior studies that have targeted mitochondria in the NPCs of the developing vertebrate brain by Khacho et al. (2017) and Iwata et al. (2020) targeted Drp1 and the control of mitochondrial structural dynamics. As a consequence, the metabolic activity of mitochondria and their control over the cell cycle were also manipulated (Mitra et al., 2009) (Madan et al., 2021). In contrast, by targeting PGC-1a we expected that rather than interfering with the function of mitochondria, we would increase or decrease organelle numbers, as has been shown previously (Rosenkranz et al., 2021).Consistent with this, we did not detect differences in mitochondria topology or structure in the NPCs (Fig. 3). Yet, by altering mitochondrial biogenesis we significantly changed how NPCs proliferated.

We boosted mitochondria levels in tectal NPCs through the overexpression of the PGC-1a (Figs 3 and 4) and limited mitochondrial biogenesis with the targeted knockdown of PGC-1a expression with the cell-targeted electroporation of a specific translation-blocking morpholino (Fig. 5). Overexpression of PGC-1a did not produce measurable changes in the distribution of mitochondria in the NPCs (Fig. 3) compared to what we observed in cells that did not express PGC-1a (Figs. 1, 2). However, when the lineages of control NPCs and those overexpressing PGC-1a were compared, we found that PGC-1a expression significantly induced neurogenic cell divisions (Fig. 4), yielding a significantly greater number of neurons over 3 days and limiting the number of NPCs. In contrast, when we knocked down PGC-1a in the tectal cells, we found that symmetric cell division was enhanced compared to the controls. Knocking down PGC-1a reduced cell proliferation overall (Fig. 5K), inhibited neurogenesis (Fig. 5J) with greater levels of symmetric cell division to increase the proportion of NPCs (Fig. 5I). In short, NPCs that overexpress PGC-1a to induce mitochondrial biogenesis prematurely are more likely to generate neurons when they divide and NPCs that are prevented from inducing mitochondrial biogenesis divide symmetrically and remained in the NPC progenitor pool. Together, our data suggest that the regulation of mitochondrial biogenesis in NPCs prior to their cell division contributes to the decision to divide symmetrically or asymmetrically.

In summary, our in vivo time-lapse imaging study provides the first images of the full structure of individual NPCs and their mitochondria before and after the cells divide. Analysis of the mitochondrial distributions in the NPCs reveals that prior to cell division, NPCs shift mitochondrial distributions toward the site of cell division, raising the intriguing possibility that like other stem cells, NPCs control the fate of their progeny by asymmetrically dividing mitochondria so that their progeny do not inherit them equally. By increasing or limiting mitochondria in NPCs through manipulations of the PGC-1a transcription factor network, our data indicate that neurogenic cell division requires a commitment to mitochondrial biogenesis before NPCs divide.

## Supporting information

Video Figure1_day1

Video Figure1_day2

Video Figure1_day13

Video Figure1_day3_timelapse

Video Figure2_day1

Video Figure3_day3

## Acknowledgements

We are grateful to the Grass Foundation for their support. We thank Adithi Ramakrishnan for her help with the radial process analysis and Dr. Jennifer Rahn for her comments on this work. Thank you to Dr. Martin K. Sekula for his valuable input and guidance on our image analysis methods.

## Competing interests

The authors declare no competing or financial interests.

## Author contributions

Conceptualization: J.E.B., Methodology: J. E. B. Validation: J.E.B.; Formal analysis: J.E.B., M.S.F. and M.R.K. Writing – original draft: J. E. B. and M. S. F.; Writing, reviewing, and editing: J. E. B. and M.R.K. Visualization: J. E. B.; Supervision: J. E. B.; Funding acquisition: J. E. B.

## Funding

This research is supported by the National Institute of Health grant HD099023 the William & Mary Charles Center for Undergraduate Research

## Notes

### Competing Interest Statement

The authors have declared no competing interest.

## REFERENCES

Agathocleous, M., Love, N.K., Randlett, O., Harris, J.J., Liu, J., Murray, A.J., Harris, W.A., 2012. Metabolic differentiation in the embryonic retina. Nat Cell Biol 14, 859–864.

Angelopoulos, I., Gakis, G., Birmpas, K., Kyrousi, C., Habeos, E.E., Kaplani, K., Lygerou, Z., Habeos, I., Taraviras, S., 2022. Metabolic regulation of the neural stem cell fate: Unraveling new connections, establishing new concepts. Front Neurosci 16, 1009125.

Aretz, I., Jakubke, C., Osman, C., 2020. Power to the daughters – mitochondrial and mtDNA transmission during cell division. Biol Chem 401, 533–546.

Arrazola, M.S., Andraini, T., Szelechowski, M., Mouledous, L., Arnaune-Pelloquin, L., Davezac, N., Belenguer, P., Rampon, C., Miquel, M.C., 2019. Mitochondria in Developmental and Adult Neurogenesis. Neurotox Res 36, 257–267.

Arshadi, C., Gunther, U., Eddison, M., Harrington, K.I.S., Ferreira, T.A., 2021. SNT: a unifying toolbox for quantification of neuronal anatomy. Nat Methods 18, 374–377.

Bahat, A., Gross, A., 2019. Mitochondrial plasticity in cell fate regulation. J Biol Chem 294, 13852–13863.

Bestman, J.E., Cline, H.T., 2020. Morpholino Studies in Xenopus Brain Development. Methods Mol Biol 2047, 377–395.

Bestman, J.E., Huang, L.C., Lee-Osbourne, J., Cheung, P., Cline, H.T., 2015. An in vivo screen to identify candidate neurogenic genes in the developing Xenopus visual system. Dev Biol 408, 269–291.

Bestman, J.E., Lee-Osbourne, J., Cline, H.T., 2012. In vivo time-lapse imaging of cell proliferation and differentiation in the optic tectum of Xenopus laevis tadpoles. J Comp Neurol 520, 401–433.

Bjornsson, C.S., Apostolopoulou, M., Tian, Y., Temple, S., 2015. It takes a village: constructing the neurogenic niche. Dev Cell 32, 435–446.

Bystron, I., Blakemore, C., Rakic, P., 2008. Development of the human cerebral cortex: Boulder Committee revisited. Nature Reviews Neuroscience 9, 110–122.

Cairns, R.A., Harris, I.S., Mak, T.W., 2011. Regulation of cancer cell metabolism. Nat Rev Cancer 11, 85–95.

Chan, D.C., 2020. Mitochondrial Dynamics and Its Involvement in Disease. Annu Rev Pathol 15, 235–259.

Chu, C.-H., Tseng, W.-W., Hsu, C.-M., Wei, A.-C., 2022. Image Analysis of the Mitochondrial Network Morphology With Applications in Cancer Research. Frontiers in Physics 10.

Collins, T.J., Bootman, M.D., 2003. Mitochondria are morphologically heterogeneous within cells. J Exp Biol 206, 1993–2000.

Cowell, R.M., Talati, P., Blake, K.R., Meador-Woodruff, J.H., Russell, J.W., 2009. Identification of novel targets for PGC-1alpha and histone deacetylase inhibitors in neuroblastoma cells. Biochem Biophys Res Commun 379, 578–582.

Dalton, C.M., Carroll, J., 2013. Biased inheritance of mitochondria during asymmetric cell division in the mouse oocyte. J Cell Sci 126, 2955–2964.

Dohla, J., Kuuluvainen, E., Gebert, N., Amaral, A., Englund, J.I., Gopalakrishnan, S., Konovalova, S., Nieminen, A.I., Salminen, E.S., Torregrosa Munumer, R., Ahlqvist, K., Yang, Y., Bui, H., Otonkoski, T., Kakela, R., Hietakangas, V., Tyynismaa, H., Ori, A., Katajisto, P., 2022. Metabolic determination of cell fate through selective inheritance of mitochondria. Nat Cell Biol 24, 148–154.

Dong, W., Lee, R.H., Xu, H., Yang, S., Pratt, K.G., Cao, V., Song, Y.K., Nurmikko, A., Aizenman, C.D., 2009. Visual avoidance in Xenopus tadpoles is correlated with the maturation of visual responses in the optic tectum. J Neurophysiol 101, 803–815.

Donnelly, M.L.L., Luke, G., Mehrotra, A., Li, X.J., Hughes, L.E., Gani, D., Ryan, M.D., 2001. Analysis of the aphthovirus 2A/2B polyprotein ‘cleavage’ mechanism indicates not a proteolytic reaction, but a novel translational effect: a putative ribosomal ‘skip’. Journal of General Virology 82, 1013–1025.

Engl, E., Attwell, D., 2015. Non-signalling energy use in the brain. J Physiol 593, 3417–3429.

Fisher, M., James-Zorn, C., Ponferrada, V., Bell, A.J., Sundararaj, N., Segerdell, E., Chaturvedi, P., Bayyari, N., Chu, S., Pells, T., Lotay, V., Agalakov, S., Wang, D.Z., Arshinoff, B.I., Foley, S., Karimi, K., Vize, P.D., Zorn, A.M., 2023. Xenbase: key features and resources of the Xenopus model organism knowledgebase. Genetics 224.

Haas, K., Jensen, K., Sin, W.C., Foa, L., Cline, H.T., 2002. Targeted electroporation in Xenopus tadpoles in vivo--from single cells to the entire brain. Differentiation 70, 148–154.

Handschin, C., Spiegelman, B.M., 2006. Peroxisome proliferator-activated receptor gamma coactivator 1 coactivators, energy homeostasis, and metabolism. Endocr Rev 27, 728–735.

Herrgen, L., Akerman, C.J., 2016. Mapping neurogenesis onset in the optic tectum of Xenopus laevis. Dev Neurobiol.

Ishihara, N., Nomura, M., Jofuku, A., Kato, H., Suzuki, S.O., Masuda, K., Otera, H., Nakanishi, Y., Nonaka, I., Goto, Y., Taguchi, N., Morinaga, H., Maeda, M., Takayanagi, R., Yokota, S., Mihara, K., 2009. Mitochondrial fission factor Drp1 is essential for embryonic development and synapse formation in mice. Nat Cell Biol 11, 958–966.

Iwata, R., Casimir, P., Vanderhaeghen, P., 2020. Mitochondrial dynamics in postmitotic cells regulate neurogenesis. Science 369, 858–862.

Katajisto, P., Dohla, J., Chaffer, C.L., Pentinmikko, N., Marjanovic, N., Iqbal, S., Zoncu, R., Chen, W., Weinberg, R.A., Sabatini, D.M., 2015. Stem cells. Asymmetric apportioning of aged mitochondria between daughter cells is required for stemness. Science 348, 340–343.

Kempermann, G., 2019. Environmental enrichment, new neurons and the neurobiology of individuality. Nat Rev Neurosci 20, 235–245.

Khacho, M., Clark, A., Svoboda, D.S., Azzi, J., MacLaurin, J.G., Meghaizel, C., Sesaki, H., Lagace, D.C., Germain, M., Harper, M.E., Park, D.S., Slack, R.S., 2016. Mitochondrial Dynamics Impacts Stem Cell Identity and Fate Decisions by Regulating a Nuclear Transcriptional Program. Cell Stem Cell 19, 232–247.

Khacho, M., Clark, A., Svoboda, D.S., MacLaurin, J.G., Lagace, D.C., Park, D.S., Slack, R.S., 2017. Mitochondrial dysfunction underlies cognitive defects as a result of neural stem cell depletion and impaired neurogenesis. Hum Mol Genet 26, 3327–3341.

Khacho, M., Harris, R., Slack, R.S., 2019. Mitochondria as central regulators of neural stem cell fate and cognitive function. Nat Rev Neurosci 20, 34–48.

Koenig, M.K., 2008. Presentation and diagnosis of mitochondrial disorders in children. Pediatr Neurol 38, 305–313.

Kosodo, Y., Toida, K., Dubreuil, V., Alexandre, P., Schenk, J., Kiyokage, E., Attardo, A., Mora-Bermudez, F., Arii, T., Clarke, J.D., Huttner, W.B., 2008. Cytokinesis of neuroepithelial cells can divide their basal process before anaphase. EMBO J 27, 3151–3163.

Kumar Sharma, R., Chafik, A., Bertolin, G., 2022. Mitochondrial transport, partitioning, and quality control at the heart of cell proliferation and fate acquisition. American journal of physiology. Cell physiology 322, C311–C325.

Lambert, T.J., 2019. FPbase: a community-editable fluorescent protein database. Nature Methods 16, 277–278.

Lau, M., Li, J., Cline, H.T., 2017. In Vivo Analysis of the Neurovascular Niche in the Developing Xenopus Brain. eNeuro 4.

Lazar, G., 1973. The development of the optic tectum in Xenopus laevis: a Golgi study. J Anat 116, 347–355.

Lee, I.W., Adhikari, D., Carroll, J., 2022. Miro1 depletion disrupts spatial distribution of mitochondria and leads to oocyte maturation defects. Front Cell Dev Biol 10, 986454.

Lin, J., Wu, P.H., Tarr, P.T., Lindenberg, K.S., St-Pierre, J., Zhang, C.Y., Mootha, V.K., Jager, S., Vianna, C.R., Reznick, R.M., Cui, L., Manieri, M., Donovan, M.X., Wu, Z., Cooper, M.P., Fan, M.C., Rohas, L.M., Zavacki, A.M., Cinti, S., Shulman, G.I., Lowell, B.B., Krainc, D., Spiegelman, B.M., 2004. Defects in adaptive energy metabolism with CNS-linked hyperactivity in PGC-1alpha null mice. Cell 119, 121–135.

Liu, K., Lin, B., Zhao, M., Yang, X., Chen, M., Gao, A., Liu, F., Que, J., Lan, X., 2013. The multiple roles for Sox2 in stem cell maintenance and tumorigenesis. Cell Signal 25, 1264–1271.

Lucas, E.K., Dougherty, S.E., McMeekin, L.J., Reid, C.S., Dobrunz, L.E., West, A.B., Hablitz, J.J., Cowell, R.M., 2014. PGC-1alpha Provides a Transcriptional Framework for Synchronous Neurotransmitter Release from Parvalbumin-Positive Interneurons. J Neurosci 34, 14375–14387.

Madan, S., Uttekar, B., Chowdhary, S., Rikhy, R., 2021. Mitochondria Lead the Way: Mitochondrial Dynamics and Function in Cellular Movements in Development and Disease. Front Cell Dev Biol 9, 781933.

Matsuda, T., Nagai, T., 2014. Quantitative measurement of intracellular protein dynamics using photobleaching or photoactivation of fluorescent proteins. Microscopy (Oxf) 63, 403–408.

Misgeld, T., Schwarz, T.L., 2017. Mitostasis in Neurons: Maintaining Mitochondria in an Extended Cellular Architecture. Neuron 96, 651–666.

Mitra, K., Wunder, C., Roysam, B., Lin, G., Lippincott-Schwartz, J., 2009. A hyperfused mitochondrial state achieved at G1-S regulates cyclin E buildup and entry into S phase. Proc Natl Acad Sci U S A 106, 11960–11965.

Miyata, T., Kawaguchi, A., Okano, H., Ogawa, M., 2001. Asymmetric inheritance of radial glial fibers by cortical neurons. Neuron 31, 727–741.

Monzel, A.S., Enriquez, J.A., Picard, M., 2023. Multifaceted mitochondria: moving mitochondrial science beyond function and dysfunction. Nat Metab 5, 546–562.

Nieuwkoop, P.D., Faber, J., 1994. Normal Table of Xenopus Laevis (Daudin): A Systematical & Chronological Survey of the Development from the Fertilized Egg Till the End of Metamorphosis, 1st ed. Garland Science, New York.

Parker, D.J., Iyer, A., Shah, S., Moran, A., Hjelmeland, A.B., Basu, M.K., Liu, R., Mitra, K., 2015. A new mitochondrial pool of cyclin E, regulated by Drp1, is linked to cell-density-dependent cell proliferation. J Cell Sci 128, 4171–4182.

Pernice, W.M., Swayne, T.C., Boldogh, I.R., Pon, L.A., 2017. Mitochondrial Tethers and Their Impact on Lifespan in Budding Yeast. Front Cell Dev Biol 5, 120.

Petridi, S., Dubal, D., Rikhy, R., van den Ameele, J., 2022. Mitochondrial respiration and dynamics of in vivo neural stem cells. Development 149.

Peunova, N., Scheinker, V., Cline, H., Enikolopov, G., 2001. Nitric Oxide Is an Essential Negative Regulator of Cell Proliferation in Xenopus Brain. J. Neurosci. 21, 8809–8818.

Pilaz, L.J., Lennox, A.L., Rouanet, J.P., Silver, D.L., 2016. Dynamic mRNA Transport and Local Translation in Radial Glial Progenitors of the Developing Brain. Curr Biol 26, 3383–3392.

Pilaz, L.J., Silver, D.L., 2017. Moving messages in the developing brain-emerging roles for mRNA transport and local translation in neural stem cells. FEBS Lett 591, 1526–1539.

Pratt, K.G., Khakhalin, A.S., 2013. Modeling human neurodevelopmental disorders in the Xenopus tadpole: from mechanisms to therapeutic targets. Disease models & mechanisms 6, 1057–1065.

Preibisch, S., Saalfeld, S., Schindelin, J., Tomancak, P., 2010. Software for bead-based registration of selective plane illumination microscopy data. Nat Methods 7, 418–419.

Rakic, P., 1972. Mode of Cell Migration to the Superficial Layers of Fetal Monkey Neocortex. J Comp Neurol 145, 61–84.

Rash, B.G., Ackman, J.B., Rakic, P., 2016. Bidirectional radial Ca(2+) activity regulates neurogenesis and migration during early cortical column formation. Sci Adv 2, e1501733.

Rash, B.G., Micali, N., Huttner, A.J., Morozov, Y.M., Horvath, T.L., Rakic, P., 2018. Metabolic regulation and glucose sensitivity of cortical radial glial cells. Proc Natl Acad Sci U S A 115, 10142–10147.

Rosenkranz, S.C., Shaposhnykov, A.A., Trager, S., Engler, J.B., Witte, M.E., Roth, V., Vieira, V., Paauw, N., Bauer, S., Schwencke-Westphal, C., Schubert, C., Bal, L.C., Schattling, B., Pless, O., van Horssen, J., Freichel, M., Friese, M.A., 2021. Enhancing mitochondrial activity in neurons protects against neurodegeneration in a mouse model of multiple sclerosis. Elife 10.

Scarpulla, R.C., 2011. Metabolic control of mitochondrial biogenesis through the PGC-1 family regulatory network. Biochim Biophys Acta 1813, 1269–1278.

Scarpulla, R.C., Vega, R.B., Kelly, D.P., 2012. Transcriptional integration of mitochondrial biogenesis. Trends in endocrinology and metabolism: TEM 23, 459–466.

Schindelin, J., Arganda-Carreras, I., Frise, E., Kaynig, V., Longair, M., Pietzsch, T., Preibisch, S., Rueden, C., Saalfeld, S., Schmid, B., Tinevez, J.Y., White, D.J., Hartenstein, V., Eliceiri, K., Tomancak, P., Cardona, A., 2012. Fiji: an open-source platform for biological-image analysis. Nat Methods 9, 676–682.

Sharma, P., Cline, H.T., 2010. Visual activity regulates neural progenitor cells in developing xenopus CNS through musashi1. Neuron 68, 442–455.

Silbereis, J.C., Pochareddy, S., Zhu, Y., Li, M., Sestan, N., 2016. The Cellular and Molecular Landscapes of the Developing Human Central Nervous System. Neuron 89, 248–268.

Sild, M., Ruthazer, E.S., 2011. Radial glia: progenitor, pathway, and partner. Neuroscientist 17, 288–302.

Sild, M., Van Horn, M.R., Schohl, A., Jia, D., Ruthazer, E.S., 2016. Neural Activity-Dependent Regulation of Radial Glial Filopodial Motility Is Mediated by Glial cGMP-Dependent Protein Kinase 1 and Contributes to Synapse Maturation in the Developing Visual System. J Neurosci 36, 5279–5288.

Silver, I.A., Erecinska, M., 1994. Extracellular glucose concentration in mammalian brain: continuous monitoring of changes during increased neuronal activy and upon limitation in oxygen supply in normo-, hypo- and hyberglycemic animals. Journal of Neuroscience 14, 5068–5076.

Spinelli, J.B., Zaganjor, E., 2022. Mitochondrial efficiency directs cell fate. Nat Cell Biol 24, 125–126.

Steib, K., Schaffner, I., Jagasia, R., Ebert, B., Lie, D.C., 2014. Mitochondria modify exercise-induced development of stem cell-derived neurons in the adult brain. J Neurosci 34, 6624–6633.

Straznicky, K., Gaze, R.M., 1972. The development of the tectum in Xenopus laevis: an autoradiographic study. J Embryol Exp Morphol 28, 87–115.

Taguchi, N., Ishihara, N., Jofuku, A., Oka, T., Mihara, K., 2007. Mitotic phosphorylation of dynamin-related GTPase Drp1 participates in mitochondrial fission. J Biol Chem 282, 11521–11529.

Tremblay, M., Fugere, V., Tsui, J., Schohl, A., Tavakoli, A., Travencolo, B.A., Costa Lda, F., Ruthazer, E.S., 2009. Regulation of radial glial motility by visual experience. J Neurosci 29, 14066–14076.

Vander Heiden, M.G., Cantley, L.C., Thompson, C.B., 2009. Understanding the Warburg effect: the metabolic requirements of cell proliferation. Science 324, 1029–1033.

Vanselow, J., Thanos, S., Godement, P., Henke-Fahle, S., Bonhoeffer, F., 1989. Spatial arrangement of radial glia and ingrowing retinal axons in the chick optic tectum during development. Brain Res Dev Brain Res 45, 15–27.

Wakabayashi, J., Zhang, Z., Wakabayashi, N., Tamura, Y., Fukaya, M., Kensler, T.W., Iijima, M., Sesaki, H., 2009. The dynamin-related GTPase Drp1 is required for embryonic and brain development in mice. J Cell Biol 186, 805–816.

Wareski, P., Vaarmann, A., Choubey, V., Safiulina, D., Liiv, J., Kuum, M., Kaasik, A., 2009. PGC-1{alpha} and PGC-1{beta} regulate mitochondrial density in neurons. J Biol Chem 284, 21379–21385.

Wehnekamp, F., Plucinska, G., Thong, R., Misgeld, T., Lamb, D.C., 2019. Nanoresolution real-time 3D orbital tracking for studying mitochondrial trafficking in vertebrate axons in vivo. Elife 8.

Weissman, T.A., Riquelme, P.A., Ivic, L., Flint, A.C., Kriegstein, A.R., 2004. Calcium waves propagate through radial glial cells and modulate proliferation in the developing neocortex. Neuron 43, 647–661.

Xavier, J.M., Rodrigues, C.M., Sola, S., 2016. Mitochondria: Major Regulators of Neural Development. Neuroscientist 22, 346–358.

Zheng, X., Boyer, L., Jin, M., Mertens, J., Kim, Y., Ma, L., Ma, L., Hamm, M., Gage, F.H., Hunter, T., 2016. Metabolic reprogramming during neuronal differentiation from aerobic glycolysis to neuronal oxidative phosphorylation. Elife 5.

